# Multimodal integration and vividness in the angular gyrus during episodic encoding and retrieval

**DOI:** 10.1101/393553

**Authors:** Roni Tibon, Delia Fuhrmann, Daniel A. Levy, Jon S. Simons, Richard Henson

**Author notes:** Corresponding author: Dr. Roni Tibon MRC Cognition & Brain Sciences Unit University of Cambridge 15 Chaucer Road Cambridge, CB2 7EF UNITED KINGDOM. The authors declare no competing financial interests.

## Abstract

Much evidence suggests that the angular gyrus (AnG) is involved in episodic memory, but its precise role is yet to be determined. We examined two possible accounts, within the same experimental paradigm: the CoBRA account (Shimamura, 2011), which suggests that the AnG acts as a convergence zone that binds multimodal episodic features; and the Subjectivity account (Yazar et al., 2012), which implicates AnG involvement in subjective mnemonic experience (such as vividness or confidence). fMRI was employed during both encoding and retrieval of paired-associates. During study, female and male human participants memorised picture-pairs of common objects (in the unimodal task) or of an object-picture and an environmental sound (in the crossmodal task). At test, they performed a cued-recall task, and further indicated the vividness of their memory. During retrieval, BOLD activation in the AnG was greatest for vividly remembered associates, consistent with the Subjectivity account. During encoding, the same effect of vividness was found, but this was further modulated by task: Greater activations were associated with subsequent recall in the crossmodal than the unimodal task. Thus, encoding data suggests an additional role to the AnG in cross-modal integration, consistent with its role at retrieval proposed by CoBRA. These results resolve some of the puzzles in the literature and indicate that the AnG can play different roles during encoding and retrieval, determined by the cognitive demands posed by different mnemonic tasks.

**Significance Statement:** We offer new insights into the multiplicity of processes that are associated with angular gyrus (AnG) activation during encoding and retrieval of newly formed memories. We used fMRI while human participants learned and subsequently recalled pairs of objects presented to the same sensory modality or to different modalities. We were able to show that the AnG is involved when vivid memories are created and retrieved, as well as when encoded information is integrated across different sensory modalities. These findings provide novel evidence for the contribution of the AnG to our subjective experience of remembering, alongside its role in integrative processes that promote subsequent memory.

## Introduction

The ventral posterior parietal cortex (vPPC), and the angular gyrus (AnG) in particular, have been associated with numerous cognitive functions, including episodic memory. Although the vPPC is considered one of the most active regions during successful episodic retrieval (reviewed by Wagner et al., 2005; Vilberg & Rugg, 2008; Shimamura, 2011; Levy, 2012; Rugg & King, 2017; Sestieri et al., 2017), patients with lateral parietal lesions can often successfully retrieve episodic memories, and are not usually considered to be amnesic.

Two main explanations have been proposed to account for AnG activation during episodic retrieval. The first derives from findings of fewer ‘remember’ responses, fewer high-confidence responses, and lack of richness, vividness and specificity of retrieved episodic events in patients with parietal lesions (Berryhill et al., 2007; Simons et al., 2010; Hower et al., 2014). Under this account (Yazar et al., 2012), vPPC involvement is viewed in terms of subjective mnemonic abilities, with the AnG involved in one’s own experience of episodic memory retrieval, rather than in objective memory performance as expressed in response accuracy. This “Subjectivity account” has been further supported by several studies with healthy individuals (Qin et al., 2011; Yazar et al., 2014; Richter et al., 2016).

The second account is the “Cortical Binding of Relational Activity” (CoBRA) theory (Shimamura, 2011), which suggests that during retrieval, the vPPC (including the AnG) acts as a convergence zone that becomes increasingly involved after initial encoding in order to bind together episodic features that are represented in disparate neocortical regions. This account builds on anatomical features of the vPPC—its central location and connections to many neocortical regions (Seghier, 2013)—making it well situated for binding episodic memories by establishing intermodal links across diverse event features. CoBRA critically predicts that tasks that demand the reinstatement of intermodal features, such as voices with faces, will depend more on the vPPC than tasks that depend on within-modality associations. This account has received support from several studies (Bonnici et al., 2016; Yazar et al., 2017), but other examinations of CoBRA were inconclusive, with multimodal integration deficits found in parietal lesion patients (Ben-Zvi et al., 2015), but no multimodal vs. unimodal associative memory effects on parietal scalp ERPs recorded in healthy volunteers (Tibon & Levy, 2014).

Even though theoretically these two accounts are distinct, in practice they are not easily dissociated, as the multisensory features comprising episodic representations enable us to experience the rich and vivid details that characterise remembering. To tease apart these accounts, we utilised a pair-associate learning paradigm, based on our prior work (Tibon & Levy, 2014; Ben-Zvi et al., 2015). fMRI of healthy volunteers was employed during two learning tasks. In the unimodal task, stimulus pairs were two semantically-unrelated object-pictures, and in the crossmodal task, stimulus pairs were an object-picture and a nameable but unrelated sound. At study, participants created a mental image of the two stimuli. At test, an object-picture was presented as the cue, and participants recalled the associated target from study, either another picture or a sound, and indicated the vividness of their memory. Thus, crossmodal “events” involved different modalities for separate objects, rather than multimodal representations of the same object. Therefore, even though these events are multimodal, they are not necessarily more vivid.

To test the retrieval-focused predictions of CoBRA and the Subjectivity accounts, we examined BOLD activation during test. The CoBRA account predicts greater parietal activity associated with objective retrieval (the difference between recall success and failure trials) in the crossmodal than the unimodal task, regardless the vividness of retrieved memories. The Subjectivity account, on the other hand, predicts a linear pattern of activity—vivid recall > non-vivid recall > recall failure—that is independent of modality. In addition, as some previous findings from patients’ studies might be due to encoding rather than retrieval deficits, we also examined memory effects from the study phase. Finally, given recent claims regarding AnG laterality effects (Bellana et al., 2016), we examined whether the left and right AnG display different memory-related activation patterns.

## Methods

### Participants

Twenty-four adults participated in the experiment (20 females, *mean age* = 25.83, *SD* = 4.02) and were reimbursed for their time. One participant who was not fully scanned due to a problem with the equipment, and two participants, who did not provide any responses in one of the experimental conditions (one with no success responses and another with no non-vivid responses), were excluded, leaving 21 participants (18 females) whose data were analysed. Participants were recruited from the volunteer panel of the MRC Cognition and Brain Science Unit in Cambridge, UK. All were fluent English speakers, MR-compatible, had normal hearing and normal or corrected-to-normal vision, and were never diagnosed with ADHD, dyslexia or any other developmental or learning disabilities. They provided informed consent for a protocol approved by a local ethics committee (Cambridge Psychological Research Ethics Committee reference PRE2016.055).

### Materials

The materials for the experiment were previously used by Tibon & Levy (2014), although some (∼10%) were replaced to improve stimulus quality. Replaced stimuli were verified as recognisable by 6 pilot participants, who did not take part in the main study. Auditory stimuli were 120 environmental sounds (such as animal sounds, tools, vehicles, etc.) downloaded from various internet sources, and edited using Audacity audio editing software, at 44 kHz, 16-bit resolution, mono mode. They were all adjusted to the same amplitude level and edited to last 2 seconds. Twelve additional sounds were used for practice trials and examples. Visual stimuli were 360 colour drawings of common objects obtained from various internet sources, including fruits and vegetables, tools, sporting goods, electrical and electronic devices, animals, food, vehicles, body parts, furniture, and clothing, each approximately 6–8 cm in on-screen size.

To form the various experimental conditions, several stimulus lists were created. Two lists of 120 pictures served as cue lists for recollection. Two additional lists served as target lists, one containing 120 pictures and the other containing 120 sounds. Each entry in the cue lists was pseudorandomly assigned to an entry in the target lists, so that all stimulus-pairs were semantically unrelated. While, unavoidably, the target lists (one list of pictures and another of sounds) remained fixed for the crossmodal and unimodal conditions, two experimental lists were created to counterbalance the cues across participants: half the participants viewed the cues from the first cue list with the auditory target and the cues from the second cue list with the visual target, and vice versa for the other half. Each experimental list was further divided into three sub-lists, to allow 1/3 of the stimuli to be presented three times (the other 2/3 presented only once, see more details below), with the sub-lists used to counter-balance the repeated stimuli across participants.

### Procedure

The paradigm used in the experiment is illustrated in Figure 1. The experiment consisted of two tasks, which took place on different days (*mean days apart* = 7.14, *SD* = 4.8). One was the crossmodal pair-associate learning and cued recall task, and the other was the unimodal pair-associate learning and cued recall task. Task order was counter-balanced across participants. On arrival at the lab, participants were seated in a quiet room, where they signed an informed consent form and performed the unscanned part of the relevant task. Participants were told that they would be presented with pairs of stimuli (sound-picture pairs in the crossmodal task and picture-picture pairs in the unimodal task), and were instructed to study those pairs for subsequent retrieval by forming an association between the stimuli. They were told that after 40 pairs had been presented, a test phase would follow, in which cue pictures alone would be presented. They would then need to recall and say the name of the paired object portrayed in the picture (in the unimodal task) or of the paired sound/object making the sound (in the crossmodal task) that had accompanied that cue. Following these instructions, a headset with a microphone was provided, together with a practice block of 5 trials. During practice, the experimenter ascertained that the participant understood the nature of the associations that were to be generated for the stimulus pairs. After completion of the practice block, participants were told that the unscanned part of the task would be repeated twice, with the same 40 pairs. They were further informed that following the unscanned part, they would be asked to perform the same task in the fMRI scanner, but for 120 pairs: 40 pairs that had just been studied and recalled at the unscanned part, and 80 new pairs. We used this two-staged procedure, where 1/3 of the stimuli are repeated thrice (twice in the unscanned part and once in the scanned part) while 2/3 of the stimuli are only shown once (in the scanned part), in order to keep our procedure as close as possible to that of Ben-Zvi et al. (2015), in which all stimuli were repeated several times, while also allowing enough recall failure trials for our contrasts of interest (pilots indicated that performance would be at ceiling if all trials were repeated more than once).

**Figure 1.**
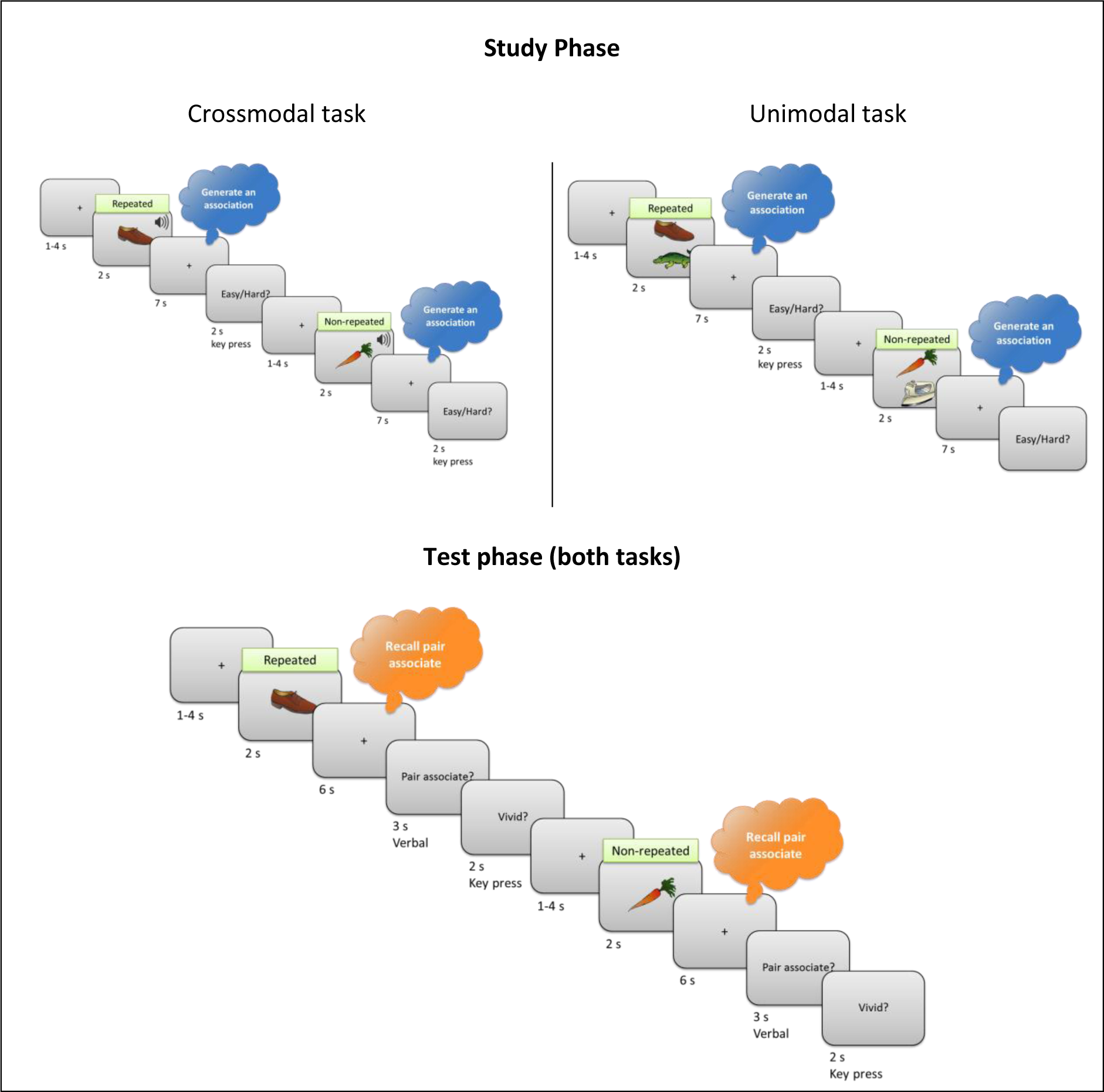
Schematic illustration of the scanned part of the experimental paradigm. During the Study phase, participants generated an association between a picture and a sound (in the crossmodal task - left) or between two pictures (in the unimodal task – right), with different tasks on different days. There were 120 stimulus pairs in total, 80 seen for the first time in the Study phase (“non-repeated”), and 40 seen twice before in a separate phase outside the scanner (“repeated”; see Methods). In the Test phase, a cue picture was presented, and participants recalled the associated sound (in the crossmodal task) or picture (in the unimodal task). Use of the same stimuli in both tasks is for illustrative purposes: in reality, each stimulus was only used in one task.

During the unscanned part of the task, participants viewed two (identical) study-test blocks of 40 stimulus pairs each. Each study trial started with a fixation cross, displayed for an exposure time jittered across a range of 1-4 s. Stimulus pairs were then presented for 2 s, followed by a 7 s fixation cross. Participants were instructed to use this time to generate an association. Next, a screen with the text “Easy/Hard?” was shown for 2 s. During this time, participants were asked to press a green key if they found it easy to come up with an association, or a red key if they found it difficult. The right index and middle fingers were used for these keypresses, and finger assignment was counterbalanced across participants.

After all 40 study trials had been presented, the test phase started. Test trials also started with a 1-4 s jittered display time fixation cross. The cue picture was then presented alone for 2 s, followed by a 6 s fixation cross. Next, a slide with the text “Pair associate?” was shown for 3 s, and participants were asked to provide their verbal response while the slide was presented. They were encouraged to formulate their response while the fixation cross was presented, so that once the “Pair associate?” slide appeared they could provide their answer immediately. They were further told that if they could not recall the target object, they should not try to guess. Instead, they should say “pass” when the “Pair associate?” slide was displayed. Finally, a slide with the legend “Vivid?” appeared for 2 s, and participants were asked to press a green key if their memory was vivid, or a red key if it was not. Finger assignment was counterbalanced across participants, and matched the assignment during the study phase (such that the same green key was used for both “Easy” and “Vivid” responses, and the same red key was used for both “Hard” and “Not vivid” responses).

After completing the unscanned part, which included 2 study-test blocks, participants were provided with instructions for the MRI scan. They were told that even though they would need to give verbal responses in the scanner, they should try to minimize head movements. They then performed the task in the MRI scanner, with one block of 120 pairs (40 pairs that were studied during the unscanned part intermixed with 80 new pairs). Finally, after the scan, participants returned to the lab where they performed a debriefing session. In this session, all target stimuli (either sounds in the crossmodal task, or pictures in the unimodal task) were displayed again, and participants were asked to type their names. The data from this session was used to subsequently eliminate (a negligible number of) trials, in which certain participants were unable to provide a name (even if inaccurate) for the sound / picture that was presented.

### fMRI Acquisition

The same acquisition and pre-processing protocol was used for both visits (one visit for the crossmodal task, and another for the unimodal task). MRI data were collected using a Siemens 3 T TIM TRIO system (Siemens, Erlangen, Germany). Structural images were acquired with a T1-weighted 3D Magnetization Prepared RApid Gradient Echo (MPRAGE) sequence [repetition time (TR) = 2250 ms; echo time (TE) = 3.02 ms; inversion time (TI) = 900 ms; 230 Hz per pixel; flip angle = 9°; field of view (FOV) 256 × 256 × 192 mm; GRAPPA acceleration factor 2]. Functional images were acquired using an echoplanar imaging (EPI) sequence with multi-band (factor = 4) acquisition. Volumes were acquired over 2 runs, one for the study phase (*mean number of volumes* = 1095.5, *SD* = 11.1) and one for the test phase (*M* = 1263, *SD* = 6.8). Each volume contained 76 slices (acquired in interleaved order within each excitation band), with a slice thickness of 2 mm and no interslice gap (TR = 1448 ms; TE = 33.4 ms; flip angle = 74°; FOV = 192 mm × 192 mm; voxel-size = 2 mm × 2 mm × 2 mm). Field maps for EPI distortion correction were also collected (TR = 541 ms; TE = 4.92 ms; flip angle = 60°; FOV = 192 × 192 mm).

### Data Pre-processing

Data were pre-processed using SPM12 (www.fil.ion.ucl.ac.uk/spm), automated in Matlab (v8.0.0.783 R2012b, The MathWorks) with Automatic Analysis (AA) 5.0 (Cusack, Vicente-Grabovetsky, Mitchell, Wild, Auer, Linke, & Peelle, 2015; https://github.com/rhodricusack/automaticanalysis). T1 anatomical images, for each participant in each visit, were coregistered to the Montreal Neurological Institute (MNI) template using rigid-body transformation, bias-corrected, and segmented by tissue class. Diffeomorphic registration was then applied across participants, separately for each visit, to the grey matter to create a group template using DARTEL (Ashburner, 2007), which was, in turn, affine-transformed to MNI space. EPI distortions in the functional images were corrected using field maps. Next, the images were corrected for motion and then for slice acquisition times by interpolating to the 26^th^ slice in time. The images were rigid-body co-registered to the corresponding T1 image and transformed to MNI space using the diffeomorphic + affine transformations. These normalised images were then smoothed by 6mm FWHM.

### Experimental Design and Statistical Analysis

#### Trial Classification

Study and test trials were classified into three response types of interest, based on subsequent response to that trial (for the study phase, to allow examination of subsequent memory effects) or actual response (for the test phase, to allow examination of recall success effects): (1) vivid: trials with correctly recalled targets, marked as “vivid”; (2) non-vivid: correctly recalled targets, classified as “non-vivid”; (3) failure: all trials with a “pass” response. Finally, trials were classified according to repetition (i.e., whether or not they were shown in the unscanned part of the task): (1) repeated: trials with correctly recalled targets, which were also shown during the unscanned part; (2) non-repeated: trials with correctly recalled targets, which were shown for the first time during the scanned part; (3) failure: all trials with a “pass” response.

The trials classification was corroborated by data obtained in the debriefing session. In cases where the participants’ response did not match the name of the object, debriefing was used to determine the source of the mismatch. The first possible source of misidentification was that the participant erroneously identified the object during the study phase (e.g., an electrical buzz sounding like a buzzing bee), and then provided the same response at debriefing and test, but that response did not match the correct response. Importantly, in this case, although the object was misidentified, memory of the target stimuli was intact. We therefore classified the trial as a success trial (either “vivid” or “non-vivid”). The second possible source of a mismatch is memory failure, i.e., the participant failed to retrieve the target, and retrieved a different object instead, therefore providing a different response at debriefing and test. In this case, the trial was classified as a “false alarm”. False alarm trials, trials in which no response was collected during the scanning session and trials in which the target was not named during the debriefing session, were marked as “excluded” (see fMRI acquisition).

#### Behavioural Analyses

All statistical analyses were performed in R v3.4.1 and RStudio v1.0153. After classifying the trials according to their responses, we analysed the behavioural data from study and test. To test for task effects (differences between the unimodal and the crossmodal task) during the study phase, data were analysed using a paired-sample t-test, to compare encoding difficulty (calculated as % of pairs for which an “easy” response was recorded at encoding) between the unimodal and the crossmodal tasks. To test for memory effects at test, accuracy data (number of trials for which vivid, non-vivid and failure responses were obtained) were analysed with a repeated measures ANOVA, which included task (crossmodal, unimodal) and response type (vivid, non-vivid, failure) as within-subject factors, using the ez package (Lawrence, 2016). Whenever sphericity assumptions were violated, Greenhouse-Geisser corrected p-values are reported. Main effects and interactions were decomposed with Bonferroni corrected pairwise comparisons using the lsmeans package (Lenth, 2016). Degrees of freedoms were corrected using the Satterthwaite method, as implemented in the lmerTest package (Kuznetsova et al., 2017).

#### ROI Analyses

To analyse ROI data, we used a mixed-models analysis, which accommodates both within- and between-subject variability. This approach is particularly recommended for unbalanced data (an unequal number of trials in each condition, Tibon & Levy, 2015), which we had here due to the post-hoc division of trials into vivid, non-vivid and failure trials (See Tibon et al., 2014; Tibon & Levy, 2014a; Tibon & Levy, 2014b for a similar use of this approach). Importantly, rather than averaged estimates across participant/condition, the mixed-models employed here require estimation of the BOLD response for each trial.

To get this estimation, we used a-priori ROIs in the AnG to extract timeseriers data. These ROIs were defined based on the coordinates of peak left AnG activation from a previous univariate meta-analysis of episodic memory by Vilberg and Rugg (2008; [−43, −66, 38]), and its homologous location in the right hemisphere [43, −66, 38]. These coordinates were also used more recently in a study by Bonnici et al. (2016). For each participant, we extracted timeseries data from these ROIs by taking the first eigenvariate across voxels within a 6mm-radius sphere centred on these coordinates, and removing effects of no interest, such as those captured the motion regressors (see whole-brain univariate analysis below). The first eigenvariate captures the dominant timeseries across voxels in an ROI, without assuming that all voxels express that timeseries equally (as assumed when simply averaging across voxels within an ROI).

To estimate the BOLD response for each trial in the extracted timeseries (separately for study and test) we used the Least Squares Separate (LSS-N) approach (Mumford et al., 2012; Abdulrahman & Henson, 2016), where N is the number of conditions. LSS-N fits a separate General Linear Model (GLM) for each trial, with one regressor for the trial of interest, and one regressor for all other trials of each condition. This implements a form of temporal smoothness regularisation on the parameter estimation (Abdulrahman & Henson, 2016). The regressors were created by convolving a delta function at the onset of each stimulus with a canonical HRF. The parameters for the regressor of interest for the ROI were then estimated using ordinary least squares, and the whole process repeated for each separate trial.

The resulting Betas were submitted to a linear mixed-model. The model included phase (study, test), task (crossmodal, unimodal), response type (vivid, non-vivid, failure), laterality (left, right) and all possible interactions between these factors as the fixed part of the model, and subject-specific slopes for each factor (see discussion by Barr et al., 2013 and Bates et al., 2018 on estimation and convergence problems of the maximal random effects model), plus a subject-specific intercept, as the random part of the model. We fitted the model using restricted maximum likelihood in R package lme4 (Kuznetsova et al., 2014) with the following formula (‘*’ indicates all possible interactions, and ‘(…|x)’ indicates random effects):

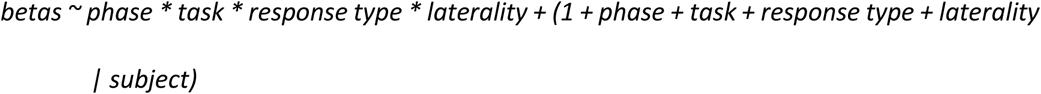

To test the predictions of the CoBRA and the Subjective accounts, we set two pre-defined contrasts, which we applied separately for each phase using the phia R package (De Rosario-Martinez, 2015). The first contrast was set to test CoBRA’s prediction of a greater objective recall success effect (success [collapsed across vividness] – failure) for crossmodal than for unimodal memories. The second contrast was set to test the prediction of the Subjectivity account of a linear pattern that is independent of modality (vivid > non-vivid > failure). Reported results are Bonferroni-corrected for two multiple comparisons (across study and test).

#### Whole-brain Univariate Analyses

We further conducted whole-brain analyses to ensure that we replicated the basic encoding and retrieval mnemonic effects found in previous studies. SPM12 and AA were used to construct General Linear Models (GLMs) for each participant, separately for each run (study, test) and for each task (crossmodal, unimodal; which took place at different visits). These first-level GLMs included 3 separate regressors for each response type of interest (vivid, non-vivid, failure) and a regressor for excluded responses (either false alarms or unnameable targets, see above). These regressors were modelled at the onset of stimulus presentation. For the study run, the GLM further included a regressor for the motor response to the “Easy/Hard” slide (locked to the motor response). For the test run, the GLM further included a regressor for the motor response to the “Vivid?” slide (locked to the motor response) and a regressor for the verbal response (at the onset of the “Pair-associate?” slide). Each of these regressors was generated with a delta function convolved with a canonical hemodynamic response function (HRF). Six subject-specific movement parameters were also included to capture residual movement-related artefacts. The GLM was fit to the data in each voxel. The autocorrelation of the error was estimated using an AR(1)-plus-white-noise model, together with a set of cosines that functioned to high-pass the model and data to 1/128 Hz (implemented to detrend the BOLD signal, and to correct physiological noises), fit using Restricted Maximum Likelihood (ReML). The estimated error autocorrelation was then used to “pre-whiten” the model and data, and ordinary least squares used to estimate the model parameters.

The images for the parameter estimates for each of the three response types for each task and each phase were then entered into second-level GLM corresponding to a repeated-measures ANOVA, which treated subject as a random effect, and with phase (study, test), task (unimodal, crossmodal) and response type (vivid, non-vivid, failure) as repeated factors. SPMs were created of the T-statistic for the “recall success” contrast of interest (success [vivid + non-vivid] – failure) at study and test. The statistical threshold was set to *p < .05* (FWE corrected for multiple comparisons across the whole brain) at the cluster-level, when the voxel-level was thresholded at *p < .0001* (uncorrected).

## Results

### Behavioural results

#### Study Phase

A paired-sample t-test revealed a significant difference in task difficulty during study, *t*(20) = 3.63, *p* = .0017, CI [.03 .13], with greater proportion of easily formed associations in the unimodal task (*M* = .69, *SD* = .19) than in the crossmodal task (*M* = .61, *SD* = .23). This difference was accounted for in our subsequent ROI analyses (see below).

#### Test Phase

The number of responses for each response type in the crossmodal and unimodal tasks are shown in Figure 2. Analysis of these data showed a significant main effect of task, *F*(1, 20) = 31.89, *p* < .0001, *η*^*2*^*G* = .01, a significant main effect of response type, *F*(2, 40) = 46.48, *p* < .0001, *η*^*2*^*G* = .65, and a significant interaction between these factors, *F*(2, 40) = 5.11, *p* = .01, *η*^*2*^*G* = .04. Bonferroni corrected pairwise comparisons showed greater number of vivid responses in the unimodal than in the crossmodal task, *t*(42.67) = 4.05, *p* = .0002, but no difference between the tasks in the amount of non-vivid or failure responses. These results support our expectation that, when different modalities are associated with separate objects within the event, rather than with the same object, crossmodal events are not necessarily remembered more vividly than unimodal events.

**Figure 2.**
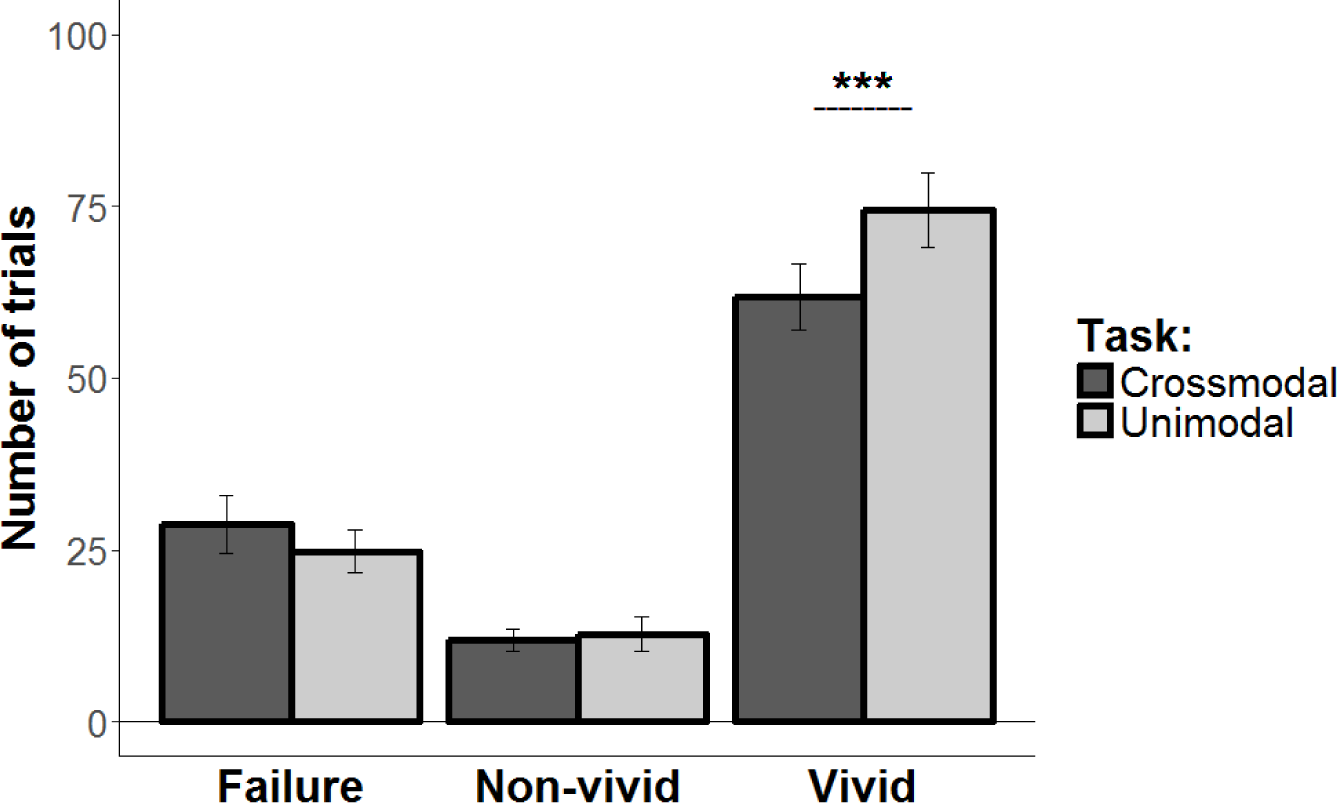
Number of responses for each response type in the crossmodal task (dark grey) and the unimodal task (light grey). Error bars represent SEs for each condition separately (*** *p* < .001).

#### ROI Mixed-Effects Analyses

For the main hypotheses under investigation, we focused on left and right AnG ROIs, defined a priori from independent data. Figure 3 depicts contrasts of the Beta values, according to our a priori contrasts of interest, averaged across left and right AnG; Table 1 shows the Betas for each condition and each ROI separately.

**Table 1.** Adjusted mean Beta values in the left and right AnG during study and test for each response type (failure, non-vivid, vivid) in each task (crossmodal, unimodal). SEs calculated for each condition separately, are given in parentheses.

|  | Left AnG |  |  | Right AnG |  |  |
| --- | --- | --- | --- | --- | --- | --- |
|  | Failure | Non-vivid | Vivid | Failure | Non-vivid | Vivid |
| *Study* |  |  |  |  |  |  |
| Crossmodal | -0.043<br>(0.007) | -0.028<br>(0.006) | -0.019<br>(0.004) | -0.037<br>(0.006) | -0.024<br>(0.006) | -0.014<br>(0.004) |
| Unimodal | -0.033<br>(0.008) | -0.034<br>(0.007) | -0.022<br>(0.006) | -0.023<br>(0.007) | -0.018<br>(0.007) | -0.009<br>(0.004) |
| *Test* |  |  |  |  |  |  |
| Crossmodal | -0.036<br>(0.007) | -0.014<br>(0.007) | -0.01<br>(0.005) | -0.035<br>(0.007) | -0.032<br>(0.007) | -0.01<br>(0.005) |
| Unimodal | -0.037<br>(0.008) | -0.026<br>(0.007) | -0.008<br>(0.006) | -0.038<br>(0.008) | -0.025<br>(0.007) | -0.005<br>(0.005) |

**Figure 3.**
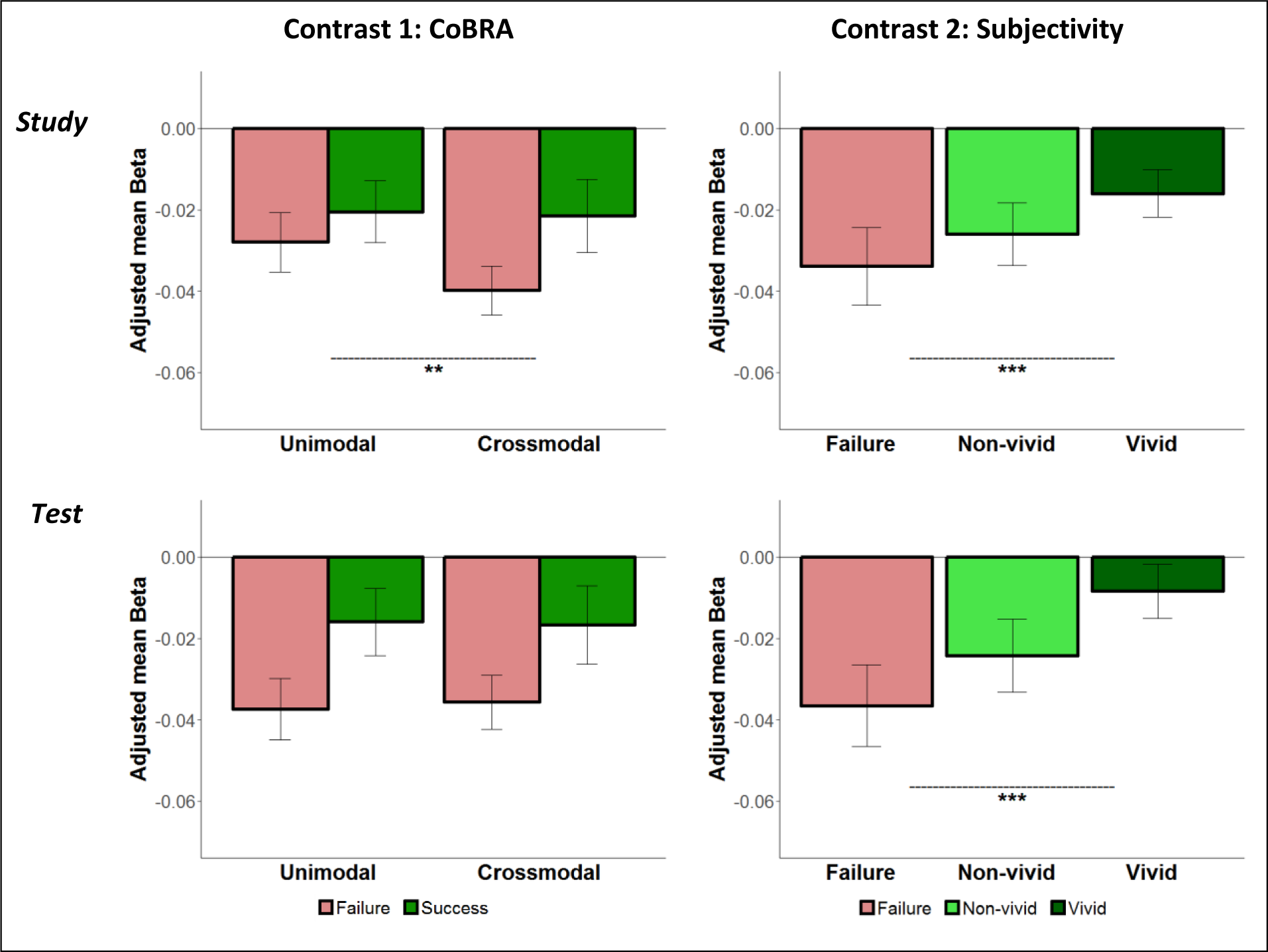
Contrasts of Beta values of the fitted model, in the AnG (collapsed across left and right AnG). Each plot represents a planned contrast: the CoBRA account (left; greater recall success effect for crossmodal than unimodal) and the Subjectivity account (right; linear trend of vivid > non-vivid > fail) during study (top panel) and test (bottom panel). Error bars represent Satterthwaite approximation of the pooled SE, and were computed for each condition separately. ** *p* < .01; *** *p* < .005.

The estimated model showed a significant main effect of response type, *χ*^*2*^(2) = 64.2, *p* < .0001, a significant interaction between phase and response type, *χ*^*2*^*(2)* = 18.1, *p* = .0001, a significant interaction between phase and laterality, *χ*^*2*^(1) = 21.33, *p* < .0001, a significant interaction between task and laterality, *χ*^*2*^(1) = 8.14, *p* = .004, and a significant 3-way interaction between phase, task, and response type, *χ*^*2*^*(2)* = 11.27, *p* = .004. Because laterality did not interact with response type, *χ*^*2*^*(2)* = 1.76, *p* = .42, we collapsed across this factor when computing our planned comparisons.

The first planned comparison confirmed a greater recall success effect for crossmodal than unimodal memories during the study phase, *χ*^*2*^*(1)* = 8.77, *p* = .006, but not during the test phase, *χ*^*2*^*(1)* = 0.19, *p* = 1. The second planned comparison confirmed a linear trend, whereby vivid > non-vivid > fail, during both study, *χ*^*2*^*(1)* = 29.14, *p* < .0001, and test, *χ*^*2*^*(1)* = 72.07, *p* < .0001.

The following analyses reported in this section were conducted to further explore these results and to rule out alternative explanations. First, we examined whether these results are also observable in the broader AnG, beyond the 6mm sphere based on the functional activation of Vilberg and Rugg (2008). To this end, we repeated the analysis on a larger, anatomical ROI from the Anatomical Automatic Labelling (AAL) atlas. This analysis revealed the same pattern as before: a significant 3-way interaction between phase, task, and response type, *χ*^*2*^*(2)* = 10.65, *p* = .005, coupled with a greater recall success effect for crossmodal vs. unimodal pairs at study, *χ*^*2*^*(1)* = 5.19, *p* = .045, but not at test, *χ*^*2*^*(1)* = 0.34, *p* = 1, and a linear trend during both study, *χ*^*2*^*(1)* = 35.98, *p* < .001, and test, *χ*^*2*^*(1)* = 102.28, *p* < .001.

Second, to ensure that the interaction between modality and response type at study cannot be explained by difficulty in forming associations (as our behavioural analysis indicated greater difficulty in the crossmodal vs. the unimodal task), we ran an additional analysis, which included the difficulty rating provided for each trial during study as a covariate in our model. Despite the addition of the covariate, the three-way interaction remained significant, *χ*^*2*^*(2)* = 11.28, *p* = .004, and the same pattern as before was confirmed by our planned comparisons, i.e., a greater recall success effect for crossmodal than unimodal memories at study phase, *χ*^*2*^*(1)* = 8.22, *p* = .008, but not at test, *χ*^*2*^*(1)* = 0.58, *p* = 09, as well as a linear trend during both study, *χ*^*2*^*(1)* = 25.46, *p* < .0001, and test, *χ*^*2*^*(1)* = 65.28, *p* < .0001.

Third, as was mentioned before, in the current experiment some of the trials were repeated several times, while others were only shown once. This was done in order to keep the procedure similar to that of Ben-Zvi et al. (2015), while also allowing for a sufficient number of failure trials. This particular design raises the possibility that the reported effects are driven by repetition, rather than by the proposed cognitive processes (i.e., multimodal integration and vividness). To address this potential confound, we ran the model again after excluding trials that were also shown during the unscanned part of the task, thus limiting our analysis to the 2/3 of the trials that were only presented once. Importantly, this analysis confirmed our previous results: the three-way interaction remained significant, *χ*^*2*^*(2)* = 9.36, *p* = .009, a greater recall success effect for crossmodal than unimodal memories was observed at the study phase, *χ*^*2*^*(1)* = 8.24, *p* = .008, but not at test, *χ*^*2*^*(1)* = 0.12, *p* = 1, and a linear trend was observed during both study, *χ*^*2*^*(1)* = 12.58, *p* < .001, and test, *χ*^*2*^*(1)* = 67.45, *p* < .0001. This indicates that repetition cannot account for the effects observed in the current study.

Finally, the repetition embedded in current design provides an opportunity to explore repetition effects in the AnG, thus tapping into CoBRA’s suggestion of greater vPPC involvement in episodic binding with the passage of time. One drawback of our design in addressing this issue is that repetition and vividness are highly correlated (inevitably, memory for repeated stimuli tends to be more vivid than memory for stimuli that were only experienced once). Nevertheless, we still sought to examine whether there are any residual repetition effects that cannot be explained by vividness. Given that the first study episode of repeated items occurred earlier than the first study episode of non-repeated ones, then according to the view that consolidation can begin immediately following initial encoding (e.g., Liu, Grady & Moscovitch, 2018), CoBRA would predict greater AnG activation for repeated vs. non-repeated items. To explore this potential effect, we added a “repetition” factor to our model, and ran a model with repetition (non-repeated, repeated), response type (vivid, non-vivid), phase (study, test), task (crossmodal, unimodal), laterality (left, right), and all possible interactions between these factors as the fixed part of the model, and with subject-specific intercept and slope for each factor as the random part of the model (because “failure” trials overlap for the “vividness” and the “repetition” factor, these trials had to be excluded in order to achieve model convergence). In general, as predicted by CoBRA, repeated trials were associated with greater AnG activation than non-repeated trials, *χ*^*2*^*(1)* = 34.15, *p* < .0001. However, the repetition effect interacted with other factors in our design, including a 3-way interaction between phase, repetition and lateralization, *χ*^*2*^*(1)* = 10.51, *p* = .001, and a 4-way interaction between repetition, response type, phase and task, *χ*^*2*^*(1)* = 5.03, *p* = .024. Given that the latter interaction is of theoretical interest, a contrast of repeated versus non-repeated trials was explored separately for each response type, phase and task. During study, this revealed a significant repetition effect for non-vivid and vivid crossmodal memories (*χ*^*2*^*(1)* = 11.99, *p* = .004; *χ*^*2*^*(1)* = 20.5, *p* < .0001, respectively) and for vivid unimodal memories (*χ*^*2*^*(1)* = 43.6, *p* < .0001). During test, this analysis revealed a significant repetition effect for non-vivid and vivid unimodal memories (*χ*^*2*^*(1)* = 20.01, *p* < .0001; *χ*^*2*^*(1)* = 9.87, *p* = .013, respectively).

#### Whole-brain Univariate Analyses

Looking beyond the AnG, to confirm that our paradigm replicated the typical engagement of regions across the brain during episodic memory encoding and retrieval, we conducted a whole-brain analysis (results are shown in Figure 4). We examined the activation to successful memory trials (collapsed across vividness), relative to failure trials, at both study and test. For both phases, this contrast yielded increased BOLD response in multiple brain regions that have previously been linked with episodic memory (see Table 2 for full results).

**Table 2.** Regions of increased BOLD activation during study and test in the recall success contrast of interest.

| Lat | Region | Peak $x,y,z$ | Cluster size | $T$ value |
| --- | --- | --- | --- | --- |
| *Study* |  |  |  |  |
| R | Precuneus | 6, -68, 46 | 1242 | 6.19 |
| R | Angular gyrus | 44, -54, 54 | 515 | 5.01 |
| R | Middle temporal gyrus | 66, -24, -8 | 80 | 5.02 |
| R | Middle cingulate gyrus | 4, -18, 30 | 214 | 4.87 |
| R | Middle frontal gyrus | 32, 66, 2 | 225 | 4.52 |
| L | Angular gyrus | -38, -62, 42 | 955 | 5.9 |
| L | Middle temporal gyrus | -64, -38, -8 | 348 | 5.63 |
| L | Middle frontal gyrus | -30, 58, 0 | 216 | 4.65 |
| *Test* |  |  |  |  |
| R | Cerebellum | 38, -56, -38 | 1376 | 7.13 |
| R | Angular gyrus | 42, -50, 20 | 1648 | 5.62 |
| R | Superior frontal gyrus | 22, 30, 48 | 482 | 5.19 |
| R | Caudate | 16, 2, 26 | 172 | 5.1 |
| R | Postcentral gyrus | 50, -10, 26 | 247 | 5.05 |
| R | Hippocampus | 28, -34, -4 | 74 | 4.99 |
| R | Middle frontal gyrus | 58, -40, -14 | 98 | 4.7 |
| L | Supramarginal gyrus<sup>a</sup> | -50, -40, 42 | 5356 | 7.9 |
| L | Angular gyrus | -50, -62, 32 |  |  |
| L | Superior frontal gyrus | -20, 26, 56 | 5047 | 7.76 |
| L | Precuneus | -6, -52, 30 | 4494 | 7.69 |
| L | Inferior frontal gyrus | -54, 12, 8 | 1882 | 6.96 |
| L | Hippocampus | -26, -28, -10 | 321 | 6.57 |
| L | Supplementary motor cortex | -4, 2, 62 | 123 | 5.95 |
| L | Putamen | -12, 8, -12 | 135 | 5.88 |
| L | Thalamus | -8, -28, 8 | 84 | 5.04 |
| L | Cerebellum<sup>b</sup> | -18, -64, -26 | 75 | 4.99 |
| L |  | -36, -74, -42 | 236 | 4.89 |

All regions arising from the contrast of interest, at a threshold of $p < 0.05$ FWE cluster-level corrected, with voxel-level threshold at $p < .0001$ (uncorrected). Anatomical labels were provided by the Neuromorphometrics atlas, available via SPM (Neuromorphometrics, Inc. http://www.neuromorphometrics.com/).
<sup>a</sup> Cluster included local maxima in the supramarginal gyrus and the angular gyrus.
<sup>b</sup> Two local maxima were observed in the left cerebellum at test.

**Figure 4.**
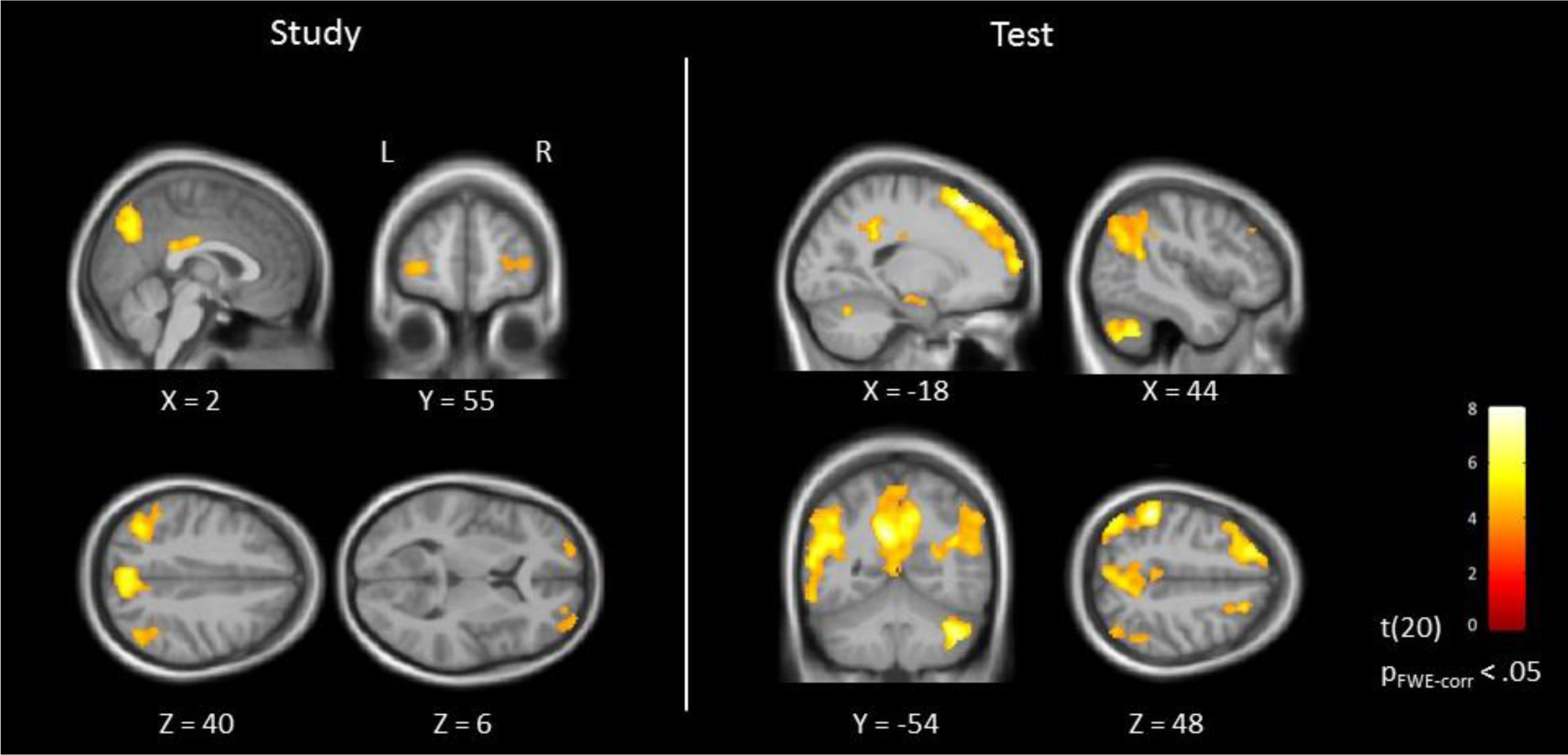
Brain regions showing stronger BOLD response for successful vs. failure recall at study (left) and test (right). *p* < .05 FWE cluster-level corrected, with voxel-level threshold at *p* < .0001 (uncorrected). Data are shown on sagittal, coronal, and axial slices of the group-averaged brain (*n* = 21).

## Discussion

The current study tested two dominant accounts of the activation in parietal cortex consistently found in neuroimaging studies of episodic memory—the CoBRA account and the Subjectivity account—and revealed a dissociation between encoding and retrieval of memory-related activations in the angular gyrus (AnG). During retrieval, we found a linear trend as predicted by the Subjectivity account: BOLD activation was greatest for vividly remembered associates and least for associates that failed to be recalled. Contrary to the prediction of the CoBRA account however, which predicts greater activation when information from multiple modalities is reinstated at retrieval, the magnitude of the objective recall success effect did not differ between the unimodal and crossmodal tasks. During encoding, activation associated with subsequent memory showed the same linear pattern predicted by the Subjectivity account, but in addition, this pattern was now moderated by task, with greater activations associated with subsequent recall success in the crossmodal than unimodal task. Memory effects at encoding and retrieval did not interact with lateralisation, suggesting that, at least with the current task, the left and right AnG play a similar role. Taken together, these results support the Subjectivity account, but not the CoBRA account as currently specified, which implicates the involvement of the AnG in multimodal reinstatement - specifically at retrieval; rather, in our current data, the involvement of AnG in multimodal processing was restricted to encoding.

Our finding that AnG activity scales with ratings of vividness during retrieval is consistent with prior studies that have found AnG to be sensitive to qualitative characteristics of memory such as vividness, confidence, and precision (e.g., Kuhl & Chun, 2014; Yazar et al., 2014; Richter et al., 2016), supporting the notion that the AnG is involved in the subjective experience of remembering. We show further here that the linear pattern associated with vivid remembering in the AnG also occurs at encoding, suggesting that it is also involved in the construction of representations that enable vivid subsequent memories. We note that this pattern of encoding-related activity, i.e., greater AnG activation for subsequently remembered vs. forgotten memories, is inconsistent with some previous studies (e.g., Daselaar et al., 2009; Lee et al., 2017), which reported negative subsequent memory effects in the AnG (that is, greater activation for subsequently forgotten vs. remembered memories). Nevertheless, an extensive meta-analysis (Uncapher & Wagner, 2009) indicated that both positive and negative subsequent memory effects are observed in the vPPC. Although a comprehensive explanation of this discrepancy is not yet available, prior literature point to two possible accounts for the positive subsequent memory effect found in the current study. First, Uncapher and Wagner (2009) suggest that retention interval is a crucial predictor of the directionality of subsequent memory effects in the vPPC, whereby positive effects are more often associated with relatively short retention times (< 45 minutes), which is consistent with our present findings. Second, Lee et al. (2017) speculate that positive subsequent memory effects would be observed in situations where the encoding of an item might benefit from retrieval and/or integration of related information (for example, when memory is measured via free recall, encoding an item as part of a broader context of temporally adjacent experiences would benefit subsequent retrieval). In the current study, participants were required to generate associations during encoding, thereby encoding the items into broader context, which might account for the positive subsequent memory effect that we have observed.

The dissociation we observe between encoding and retrieval potentially resolves a puzzle arising from our prior studies using a very similar paradigm, where we found that patients with parietal lesions did show a deficit in multimodal reinstatement (Ben-Zvi et al., 2015), prima facie supporting the CoBRA account, but we also failed to find any effect of multimodal reinstatement on parietal ERPs recorded during retrieval in healthy volunteers (Tibon & Levy, 2014), contrary to the CoBRA account. The lack of effects of multimodal reinstatement at retrieval in the present fMRI study agree with the prior ERP study, while the effect of multimodal processing that was found at encoding suggests that the deficit in patients might arise when they encode the paired-associates, rather than being a problem at retrieval.

Our finding that AnG activation is not modulated by multimodal reinstatement during retrieval needs to be considered with respect to other findings. In particular, two previous studies have shown retrieval-related involvement of the AnG in tasks that require multimodal reinstatement. In the first study, Yazar et al. (2017) used continuous post-encoding theta burst transcranial magnetic stimulation to interrupt AnG functioning. Participants encoded objects which were presented both visually and auditorily, and were subsequently asked to retrieve unimodal sources (e.g., which side + which location) or crossmodal sources (e.g., which side + male or female voice) of the studied objects. Following stimulation, participants’ ability to retrieve information was reduced for crossmodal but not for unimodal sources. In the second study, by Bonnici et al. (2016), participants memorised audio clips, visual clips, and audio-visual clips, presenting different objects (e.g., a train). At test, they were given a verbal cue for the to-be-recalled clip (e.g., the word “train”), and were asked to recall the associated clip as vividly as possible. This study showed greater AnG activation when crossmodal than unimodal memories were retrieved, coupled with above chance classification accuracy of individual crossmodal (but not unimodal) memories in the AnG. This previous evidence of AnG involvement in multimodal retrieval effects suggest that although our current results seem to dissociate between encoding and retrieval of memory-related activations in AnG, a simple distinction between memory stages might be too simplistic. Instead, we suggest that what determines AnG involvement is not the memory stage per-se (encoding vs. retrieval), but rather the specific combination of cognitive demands posed by the task. Hence, unlike CoBRA, which predicts AnG involvement whenever multimodal elements are reinstated, we suggest that such activation might be further constrained by other task characteristics, such as the type of information that is retrieved and the mnemonic cues that are provided (for example, it might be that AnG activation is only triggered when the retrieved item itself contains multimodal information, or when source/contextual information, rather than core item information, is being retrieved). This suggestion is in line with the component process model of memory (e.g., Moscovitch, 1992; Witherspoon & Moscovitch, 1989), which posits that numerous different processing components, associated with distinct brain regions, are recruited in various combinations by different memory tasks. Thus, different task demands would involve distinct process-specific alliances (that is, transient interactions between several brain regions, Cabeza & Moscovitch, 2013; Moscovitch et al., 2016). Notably, in accord with this view, even though some aspects of our results seem to be inconsistent with previous studies that used different tasks (Bonnici et al., 2016; Yazar et al., 2017), our findings do converge across the three studies that used the same paradigm (i.e., the current study, Ben-Zvi et al., 2015, and Tibon & Levy, 2014), where task demands were kept similar. Importantly, by elucidating the nature of process-specific alliances, future studies can provide a more decisive and fine-grained account of AnG involvement in multimodal processing.

In addition to the main processes investigated here, our study offers some ancillary insights regarding repetition effects in the AnG. Greater activity for repeated versus non-repeated items was consistently observed during the study phase. This finding is consistent with CoBRA’s suggestion that vPPC involvement begins after initial encoding, and increases as time passes. One exception, where this repetition effect was not significant at study, was the case of non-vivid unimodal memories. We speculate that in this case, repeated trials (subsequently classified as non-vivid) were forgotten following their initial presentation, and therefore experienced as new during the scanned part of the task. This would make them more similar to non-repeated trials, and possibly eliminate the repetition effect (why the same pattern was not observed in the crossmodal task is unclear, but might be related to a difference in the criterion set for trial classification as vivid or non-vivid in this task). During retrieval, repetition effects were observed in the unimodal task, but not in the crossmodal task. CoBRA suggests that vPPC acts as a way of offloading reliance on MTL bindings. The retrieval repetition effect might therefore indicate that responsibility for unimodal memories rapidly shifts from MTL to vPPC, whereas crossmodal memories require prolong MTL binding prior to vPPC involvement. Importantly, however, the exploration of repetition effects goes beyond the original purpose of our design, and therefore these suggested interpretations of repetition effects should be treated with caution.

In the current study, unimodal associations were comprised of two visual stimuli, and crossmodal associations were comprised of a visual and an auditory stimulus. We chose these materials to allow simultaneous presentation of the stimuli (given that simultaneous processing of two auditory stimuli, for example, is perceptually challenging). However, this means that a limitation of the current study is that we cannot be certain that our conclusions generalise to other kinds of materials. It could be, for example, that the AnG is selectively involved in multimodal integration of audio-visual associations, but not in the integration of information deriving from other sensory modalities. While we have no reason to assume that audio-visual associations are unique in this sense, future studies can use a similar paradigm to test other forms of within- and between-modality combinations. Another potential limitation of the current study is the uneven distribution of female (n=18) and male (n=3) participants. Although we do not expect the mnemonic processes addressed in this study to differ between genders, generalisation of our findings to males should be done cautiously.

In summary, the results of the current study show that the AnG is involved in multiple mnemonic processes, during both encoding and retrieval. They provide a straightforward answer to the puzzle arising from our previous findings (in Ben-Zvi et al., 2015 versus Tibon & Levy, 2014), by suggesting that the AnG is involved in (at least) two memory-related processes: multimodal integration at encoding, and construction of representations that enable vivid recall, either during encoding or during retrieval. Based on current and prior evidence (Bonnici et al., 2016; Yazar et al., 2017), it is plausible that the extent of AnG activation in mnemonic processes is determined by specific cognitive demands posed by the task (e.g., Moscovitch et al., 2016). Future studies might systematically manipulate such demands within the same experimental paradigm, and examine activation during encoding and retrieval, possibly using a more fine-grained vividness rating (see also Richter et al., 2017). Nonetheless, our current results provide an important step towards clarification of the complexities regarding AnG involvement in episodic memories.

## Acknowledgements

RT is supported by a Newton International Fellowship by the Royal Society and the British Academy (grant SUAI/009 RG91715) and by a British Academy Postdoctoral Fellowship (grant SUAI/028 RG94188). JSS is supported by the James S McDonnell Foundation (#220020333). RH is supported by UK Medical Research Council grant (SUAG/010 RG91365). The authors wish to thank Josefina Weinerova for her assistance in collecting the data and Yuval Suchowski for help with figures preparation.

The authors declare no competing financial interests.

